# The association between low-density lipoprotein cholesterol predicted by *HMGCR* genetic variants and breast cancer risk may be mediated by body mass index

**DOI:** 10.1101/464446

**Authors:** Siddhartha P. Kar, Hermann Brenner, Graham G. Giles, Dezheng Huo, Roger L. Milne, Gad Rennert, Jacques Simard, Wei Zheng, Stephen Burgess, Paul D. P. Pharoah

**Author notes:** Correspondence to Siddhartha Kar.

## Abstract

Orho-Melander et al. recently reported that lower low-density lipoprotein cholesterol (LDLC) as predicted by the T-allele of the variant rs12916 in *HMGCR* is associated with a decreased risk of developing breast cancer [odds ratio (OR) = 0.89; 95% confidence interval (CI): 0.82–0.96].^1^ This analysis was embedded in a wider Mendelian randomization (MR) study performed using genotype data from a prospective cohort of 26,589 individuals that included 16,022 women and 1176 incident breast cancer cases. *HMGCR* encodes 3-hydroxy-3-methylglutaryl-coenzyme A reductase, the enzyme inhibited by statins. The T-allele of rs12916 is associated with reduced *HMGCR* expression and therefore, in principle, its effects should be analogous to the effects of lifelong statin administration starting at birth.^2^ The MR study of Orho-Melander et al. also found that a genome-wide LDLC score based on 32 independent LDLC-associated single nucleotide polymorphisms (SNPs) was not associated with breast cancer. In light of this finding, they suggest that the protective effect of the rs12916 T-allele on breast cancer may either be specific to LDLC lowering via genetic inhibition of *HMGCR* or be the result of a distinct mechanism that is regulated by rs12916 and *HMGCR*.

---

Orho-Melander et al. reported that rs12916 was not associated with body mass index (BMI) in their cohort (*P* = 0.29). However, we noted that this SNP has previously been identified for its association with BMI in adults in a genome-wide association meta-analysis of 315,585 individuals (Table 1)^3^ This pleiotropic BMI-increasing effect of the LDLC-lowering T-allele of rs12916 is noteworthy because it has been demonstrated in another large MR study by Guo et al. that higher genetically-predicted adult BMI is associated with reduced risk of estrogen receptor (ER)-positive and ER-negative breast cancer.^4^ We hypothesized that the pleiotropic BMI-increasing effect of *HMGCR* rs12916 T-allele, rather than its LDLC-lowering effect, may be responsible for the reported association with breast cancer protection.

**Table 1:** Summary genetic association results for individual SNPs in the *HMGCR* region.^a^

| SNP | EA | LDLC |  | BMI |  | ER-pos BC |  | ER-neg BC |  | Overall BC |  |
| --- | --- | --- | --- | --- | --- | --- | --- | --- | --- | --- | --- |
|  |  | Beta | P | Beta | P | Beta | P | Beta | P | Beta | P |
| rs10066707 | G | -0.0497 | 3.0E-19 | 0.0097 | 0.02 | -0.0112 | 0.15 | -0.0113 | 0.34 | -0.0136 | 0.04 |
| rs12916 | T | -0.0733 | 7.8E-78 | 0.0182 | 7.30E-09 | -0.0136 | 0.07 | -0.0105 | 0.36 | -0.0113 | 0.07 |
| rs17238484 | G | -0.0627 | 1.4E-21 | -- | -- | -0.0237 | 0.006 | -0.0151 | 0.26 | -0.0156 | 0.03 |
| rs2006760 | C | -0.0533 | 1.7E-13 | 0.006 | 0.27 | -0.0161 | 0.08 | -0.0052 | 0.71 | -0.0135 | 0.08 |
| rs2303152 | G | -0.0423 | 1.0E-09 | 0.0158 | 2.8E-03 | -0.0222 | 0.07 | -0.0005 | 0.98 | -0.0077 | 0.45 |
| rs5909 | G | -0.0617 | 4.9E-13 | -0.0008 | 0.90 | 0.0066 | 0.59 | 0.0057 | 0.76 | -0.0029 | 0.78 |
| rs2112347<sup>b</sup> | T | -0.0443 | 4.4E-30 | 0.0261 | 6.2E-17 | -0.0154 | 0.04 | -0.0343 | 3.3E-03 | -0.0183 | 4.4E-03 |

<sup>a</sup>Obtained from the data sets in references 3, 9, and 10.
<sup>b</sup>rs2112347 is the SNP most strongly associated with BMI in the *HMGCR* region in the BMI genetic association meta-analysis.<sup>3</sup>
Abbreviations: EA: effect allele; BC: breast cancer; ER-pos: estrogen receptor positive; ER-neg: estrogen receptor negative; LDLC: low density lipoprotein cholesterol; BMI: body mass index.

We further explored this hypothesis using a two-sample Mendelian randomization study with a multivariable approach.^5,6^ It is not possible to perform a multivariable MR analysis using only one SNP as a genetic instrument and so we constructed a multi-SNP instrumental variable for this purpose. Six SNPs in or within 100 kb of *HMGCR* (including rs12916) have previously been used to construct a genetic instrument for LDLC lowering via inhibition of *HMGCR* in studies that have compared this genetically-predicted effect with the actual effect of statins on cardiovascular outcomes observed in clinical trials.^7,8^ Forward stepwise-regression models based on individual-level data have shown that each of these SNPs has a strong and independent effect on circulating LDLC level.^7^ We extracted summary results for these SNPs (Table 1) from publicly available genetic association meta-analysis data sets for LDLC (maximum n = 168,357 individuals),^9^ BMI (max. n = 315,585 individuals),^3^ estrogen receptor(ER)-positive breast cancer risk (69,501 cases/105,974 controls),^10^ ER-negative breast cancer risk (21,468 cases/105,974 controls),^10^ and overall breast cancer risk (122,977 cases/105,974 controls).^10^ The overall breast cancer risk data set included breast cancer cases whose ER-status was unknown, in addition to all ER-positive and ER-negative cases included in the two ER-specific data sets. One of the six SNPs, rs17238484, had not been mapped in the BMI genetic association meta-analysis.^3^ For four of the five SNPs with BMI data available, the LDLC-lowering allele was the BMI-increasing allele and conferred protection against breast cancer (Table 1). We used a random-effect linear regression model weighted by the inverse of the variance of the SNP associations with breast cancer for MR analysis.^11,12^ We obtained the genetic correlations between the six SNPs using the ‘TwoSampleMR’ *R* package and accounted for the effect of these genetic correlations in our MR analyses using the method implemented in the ‘MendelianRandomization’ *R* package.^12,13^ Univariable analysis confirmed that lower LDLC as predicted by all six *HMGCR* SNPs put together was associated with reduced ER-positive and overall breast cancer risk (Figure 1). For the corresponding ER-negative breast cancer risk analysis, while the confidence interval included one, the point estimate of the OR was consistent in magnitude and direction with the ER-positive and overall breast cancer results (Figure 1). We also obtained similar point estimates when using rs12916 alone as a genetic instrument to test the association between LDLC and breast cancer risk by the Wald ratio method (Figure 1).^13^ However, multivariable Mendelian randomization analysis, wherein standardized regression coefficients from the genotype-BMI association analysis were included as a covariate in the model,^6^ attenuated the protective effect on breast cancer risk of lower LDLC as collectively predicted by the five *HMGCR* SNPs that had both LDLC and BMI data available (Figure 1). The shift in OR towards the null after adjusting for the effect of BMI was marked for ER-positive breast cancer (from 0.83 to 0.99) and ER-negative breast cancer (from 0.86 to 0.97) but small for the overall breast cancer data (from 0.84 to 0.86). On balance, the multivariable MR analyses suggested that the pleiotropic effects on BMI may mediate the association between LDLC-lowering alleles in *HMGCR* and reduced breast cancer risk.

**Figure 1:**
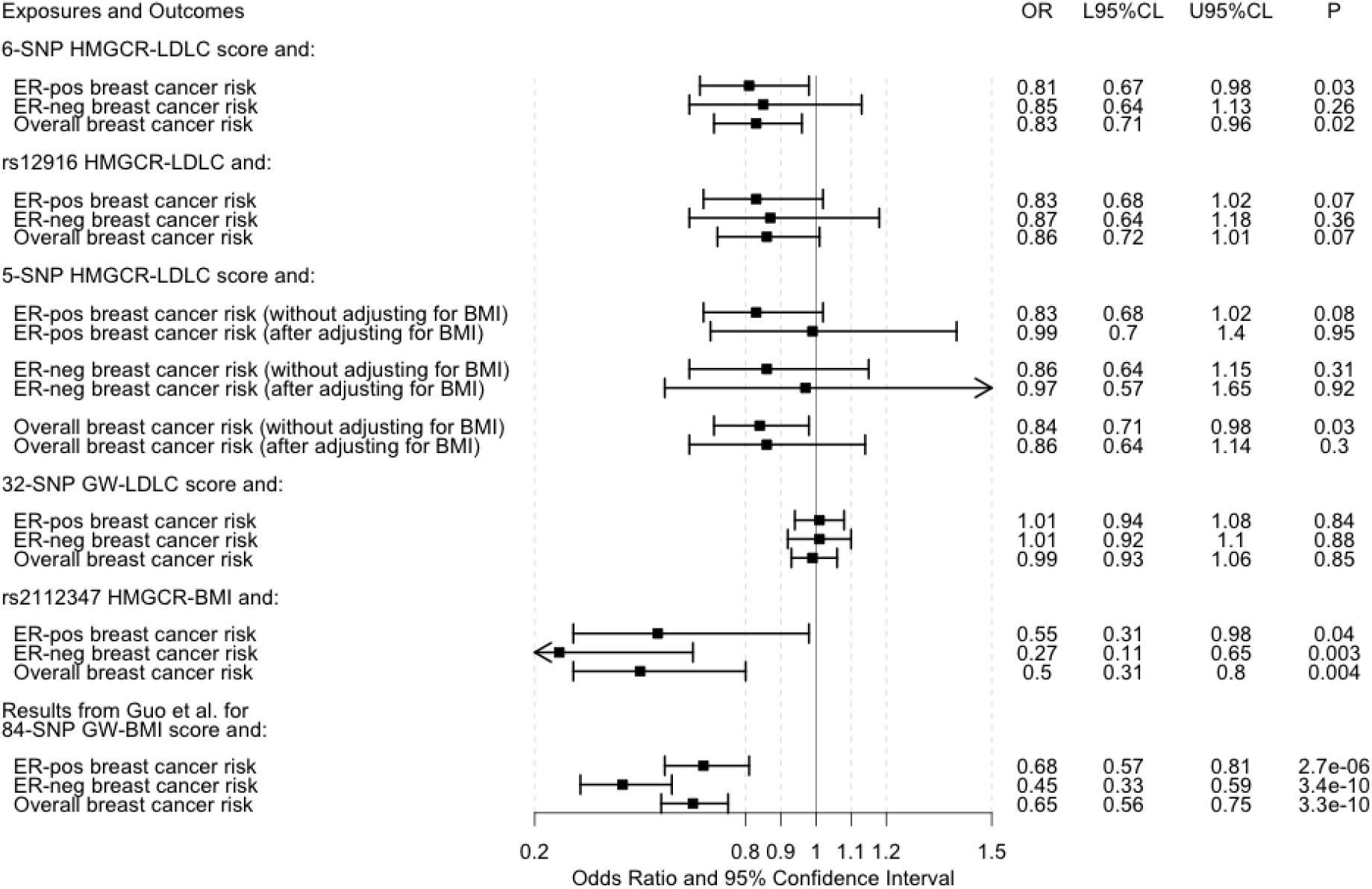
Forest plot of the results of Mendelian randomization analyses. The odds ratios (ORs) presented for breast cancer risk are per 1-standard deviation (~1 mmol/L) decrease in low-density lipoprotein cholesterol (LDLC) or per 1-standard deviation (~4.65 kg/m^2^) increase in adult body mass index (BMI) except for the ORs from Guo et al.,^4^ which are scaled per 5 kg/m^2^ increase in adult BMI. The estrogen receptor (ER)-positive (ER-pos), ER-negative (ER-neg), and overall breast cancer results from Guo et al.^4^ were based on analyses of 69,556, 49,770, and 88,807 women, respectively, and this data set overlapped entirely with the data set in reference 10. Other abbreviations: L95%CL: lower 95% confidence limit; U95%CL: upper 95% confidence limit; SNP: single nucleotide polymorphism; GW: genome-wide.

We confirmed that the 32-SNP genome-wide LDLC score from Orho-Melander et al. was not associated with breast cancer risk in our data sets (using beta coefficients for these SNPs from the 2013 Global Lipid Genetics Consortium data set^9^ and without adjustment for BMI; Figure 1). We also used the Wald ratio method to show that higher BMI predicted by the SNP that had the strongest association with BMI in the *HMGCR* genomic region,^3^ rs2112347 which is 321 kb from *HMGCR* and has a correlation (*r*^2^)of 0.37 with rs12916 in European-ancestry populations from the 1000 Genomes Project,^14^ was associated with lower risk of ER-positive, ER-negative, and overall breast cancer (Figure 1). Taken together with the inverse association between BMI genetically-predicted by 84 SNPs across the genome and breast cancer risk identified by Guo et al. (Figure 1),^4^ these findings lend additional support to BMI, rather than LDLC, being a more likely mechanism underlying the potential protective effect of genetic inhibition of *HMGCR* on breast cancer risk that was identified by Orho-Melander et al.

## Acknowledgements

This work builds on multiple, publicly available data sets. The low-density lipoprotein cholesterol association data were obtained from the Global Lipid Genetics Consortium (GLGC). The body mass index association data were obtained from the Genetic Investigation of ANthropometric Traits (GIANT) Consortium. The breast cancer genetic association data were obtained from the Breast Cancer Association Consortium (BCAC).

## Data sets

**Global Lipid Genetics Consortium (GLGC):**

http://csg.sph.umich.edu/abecasis/public/lipids2013/

**Genetic Investigation of ANthropometric Traits (GIANT) consortium:**

https://portals.broadinstitute.org/collaboration/giant/index.php/GIANT_consortium_data_files#GWAS_Anthropometric_2015_BMI

**Breast Cancer Association Consortium (BCAC):**

http://bcac.ccge.medschl.cam.ac.uk/bcacdata/oncoarray/gwas-icogs-and-oncoarray-summary-results/

